# Exact polynomial-time graph canonisation and isomorphism testing through comparison of ordered vertex eigenprojections

**DOI:** 10.1101/2022.04.14.488296

**Authors:** Robert John O’Shea

## Abstract

**Motivation:** Graph canonisation and isomorphism testing representation are fundamental computational problems, whose complexity has remained unsolved to date. This study examines graph eigenprojections, demonstrating that linear-ordering transformations induce canonical properties therein to yield polynomial-time canonisation and isomorphism testing in all undirected graphs.

**Results:** This study presents an exact method to identify analogous vertices in isomorphic graphs, through comparison of vertices’ eigenprojection matrices, which are shown to be related by a linear permutation. Systematic perturbation strategies are developed to reduce degeneracy whilst conserving isomorphism, through the addition of characteristically weighted self-loops to analogous vertices. Repeated iterations of analogy testing and perturbation deliver canonical vertex labelling and recovery of isomorphic mappings in 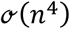 time in all graphs. Analytical proofs are provided to support claims and experimental performance is demonstrated in biological and synthetic data, with comparison to a commonly used heuristic algorithm.

**Availability and Implementation:** Source code is provided at github.com/robertoshea/graph_isomorphism.

**Supplementary Data.:** Not applicable.

## Introduction

The graph isomorphism problem requires the determination of structural equivalence of two graphs. It is considered one of the most important unresolved problems in computer science (Grohe and Schweitzer, 2020) and arises in applications across the domains of image recognition, systems biology and cheminformatics (Grohe and Schweitzer, 2020). Although the problem’s computational complexity has remained an open problem since its description in 1972 (Karp, 1972; Grohe and Schweitzer, 2020), it has been demonstrated that it is in the low hierarchy of the class NP (Schöning, 1988; Arvind and Torán, 2005). Graph canonisation is a closely related task, which requires graph representation in a form which is invariant under permutation and characteristic of the graph automorphism group. Thus, isomorphism testing may also be achieved by representing both graphs in canonical form and checking equality.

Exact polynomial time algorithms are available for some special cases of the graph isomorphism problem. Babai proposed a spectral assignment algorithm for graphs with bounded eigenvalue multiplicity (Babai *et al*., 1982). Luks proposed an algorithm for graphs of bounded degree (Luks, 1982) which was developed further by Babai to provide a quasipolynomial-time solution for the general case (Babai, 2016; Helfgott *et al*., 2017).

Klus and Sahai developed a heuristic spectral assignment algorithm for degenerate graphs, in which vertex analogy is estimated by comparing vertex eigenpolytopes (Klus and Sahai, 2018). A graph perturbation strategy was introduced in which pairs of analogous vertices were sought and perturbations applied to induce unique analogies. Analogous vertices were estimated by sorting individual rows of vertices’ eigenprojection matrices and subsequently testing equality. As the algorithm was not constrained to apply the same sorting permutation to each row, equality could be induced between some non-analogous vertex pairs, yielding false positives. Subsequently, when perturbations were erroneously applied to non-analogous vertex pairs, “backtracking” was required (Klus and Sahai, 2018). No exact polynomial-time algorithm has been identified for the general case graph isomorphism problem to date (Grohe and Schweitzer, 2020; Klus and Sahai, 2018).

## Hypothesis and Objectives

It was hypothesised that Klus’ heuristic could be developed into an exact method by modifying their analogy testing and perturbation strategies. It was hypothesised that a linear permutation would induce equality between analogous vertices’ eigenprojection matrices, which would suffice to demonstrate analogy. The objective of this study is to prove this result and employ it to develop exact polynomial-time algorithms for graph canonisation and isomorphism testing. Accordingly, an exact method is developed to identify analogous vertex pairs through comparison of vertices’ eigenprojection matrices. Klus’ perturbation strategy is developed to reduce degeneracy in a canonically reproducible manner, conserving isomorphism of perturbed graph tuples by perturbing pairs of analogous vertices. Theoretical analysis is provided to support these claims and experimental performance is demonstrated in real and synthetic data.

## Systems and Methods

### Preliminaries and notation

Let 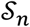 denote the symmetric group of degree *n* and 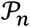 denote the set of *n* × *n* dimensional permutation. Let 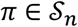 and 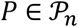 denote corresponding vector and matrix representations of the same permutation, such that:

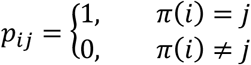

Throughout the article, vector and matrix representation of permutation functions will be used interchangeably.

Let 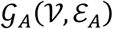 and 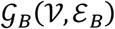 be two weighted undirected graphs, with adjacency matrices 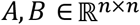, where 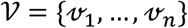 is the vertex set and *ε_A_* and *ε_B_* are edge sets.

#### Definition.

Graphs 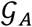 and 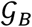 are termed “isomorphic” if there exists some permutation 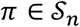 such that:

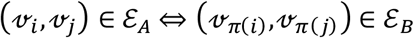

Equivalently, 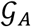 and 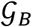 are isomorphic if there exists some 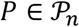, such that:

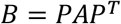

Isomorphism of 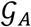 and 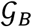 is denoted 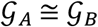.

#### Definition.

A permutation 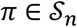 is termed an “isomorphic mapping” from 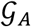 to 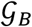 if it satisfies:

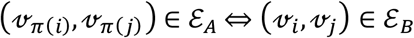

Equivalently, 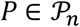 is termed an isomorphic mapping from 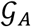 to 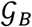 if it satisfies *B* = *PAP^T^*. This is denoted 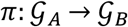. Likewise, a permutation 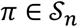 is termed an isomorphic mapping from 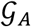 to 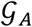 if it satisfies:

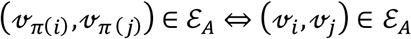

It is emphasised here that the double right arrow “⇒” will be used to denote implication of the right statement by the left., which is distinct from the single left-right arrow “→” used to denote function mappings.

#### Definition.

“Analogy” of the *i*th vertex in 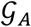 to the *j*th vertex in 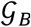, implies that there exists some 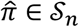 such that 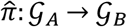 and 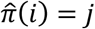. This is denoted 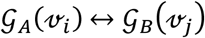. Likewise, 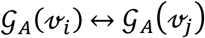 implies that there exists some 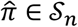 such that 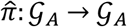 and 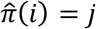. It is emphasised here that the double left-right arrow “⇔” will be used to denote equivalence of statements, which is distinct from the single left-right arrow “↔” used to denote analogy.

#### Definition.

A “unique analogy” of the *i*th vertex in 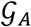 to the *j*th vertex in 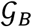 implies that:

If *x* = *i* and *y* = *j* then:

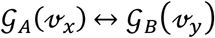

Otherwise:

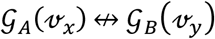

Unique analogy of the *i*th vertex in 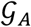 to the *j*th vertex in 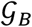 is denoted 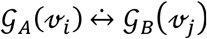.

#### Definition.

The “automorphism group” of 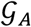, denoted 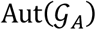 is the set of all graphs 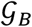 satisfying 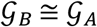.

#### Definition.

A “canonical labelling function” is a function mapping 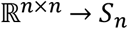 such that for any 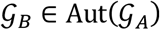, labellings *γ_A_* and *γ_B_* satisfy:

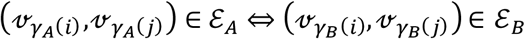

Let 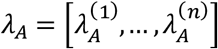 denote the eigenvalues of 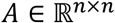 such that 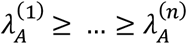. Let 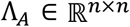 denote diag(*λ_A_*).

Let 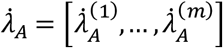 denote the *m* ≤ *n* distinct eigenvalues of *A*, such that 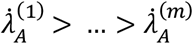.

Let 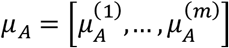 denote the eigenvalue multiplicities of *A*, such that:

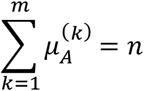

Let 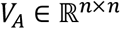 be an eigenbasis of *A*. Let *V_A_* be partitioned by eigenvalue, such that 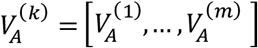, where 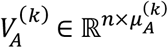.

#### Definition.

Let 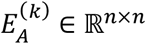 be the orthogonal projection of *A* onto its *k*th eigenspace, such that:

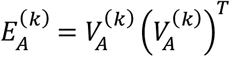

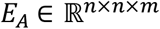 is termed the “tensor of eigenspace projections”, such that 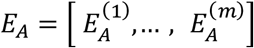.

Thus, we have:

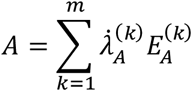

#### Definition.

Let 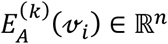 denote the *i*th column of 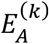, representing the projection of the *i*th vertex onto the fcth eigenspace. The matrix of a vertex’s projections onto each eigenspace is termed the “vertex eigenprojection”. 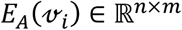 denotes the *i*th vertex eigenprojection in *A*, such that:

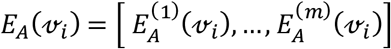

#### Definition.

For some matrix 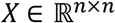, let 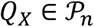 be a row-permutation matrix satisfying:

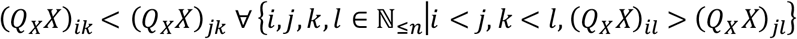

Thus, pre-multiplication by *Q_X_* orders rows in ascension by the first column, then the second, then the third etc, resolving ties in former columns by elements of latter columns. It is noted that *Q_X_* may have multiple solutions if some rows *X* are identical. Let 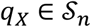 denote the permutation vector corresponding to *Q_X_*.

#### Definition.

Let 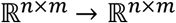 denote the row ordering function *s*(*X*) = *Q_X_X*.

#### Definition.

The row-permutation of a vertex eigenprojection under *s* is termed the “ordered vertex eigenprojection”. The *i*th ordered vertex eigenprojection in *A* is given by 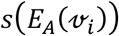.

#### Definition.

Let 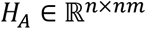 denote the matrix of ordered vertex eigenprojections in vectorised form, such that the *i*th row, 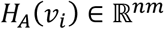, is given by:

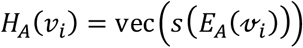

Let *I_n_* denote the *n*-dimensional identity matrix. Let 1*_n_* denote an *n* × 1 column of ones and let 1*_n_*×*_n_* denote a matrix of ones.

#### Definition.

Let 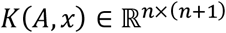 denote the Krylov matrix (Dax, 2017) generated by the product of 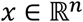 and exponents of 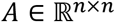 such that

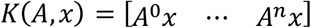

### Algorithm Overview

This study presents an exact method to identify analogous vertices in isomorphic graphs, through modification of Klus’ algorithm. It is demonstrated that the ordered eigenprojections of analogous vertices are equal. Furthermore, it is demonstrated that equality of ordered vertex eigenprojections implies analogy of vertices. These results may be exploited to test analogy of any vertex pair. An isomorphism-preserving perturbation strategy is introduced, in which pairs of analogous vertices are perturbed with characteristically weighted self-loops, inducing uniqueness of their analogy. The steps of analogy-testing and perturbation may be iterated until all analogies in the perturbed graphs are unique. It is demonstrated that fewer than *n* iterations are required to infer an isomorphic mapping in this way. Thus, isomorphic mappings may be extracted in 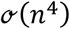 time. Two algorithms are presented to infer mappings between isomorphic graph tuples. In the first algorithm, graphs are perturbed simultaneously. In the second algorithm, the procedure is optimised by iteratively generating canonical orderings for each graph, reducing the number of vertex analogy tests required.

### Analogy Testing

When searching for an isomorphic mapping between two graphs, it is appropriate to presume graphs are isomorphic, and test this assumption when a candidate isomorphic mapping has been identified. Accordingly, we let 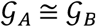. **Lemma 1** demonstrates that *E_A_* is invariant to choice of eigenbasis, even when spectra are degenerate.

#### Lemma 1.

If *B* = *PAP^T^* then 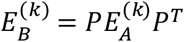 for all *k* ∈ {1,&, *m*}.

*Proof*

We have:

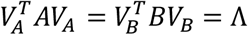

As Λ is diagonal, it may be decomposed by eigenspace, such that:

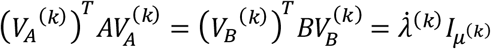

Substituting *B* = *PAP^T^*, we have

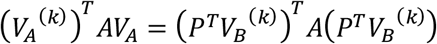

Removing *A*, we have:

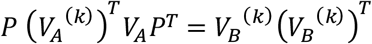

Substituting the definition of 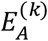 and 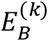 we have:

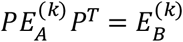

***End of lemma 1***

Hence, the projections 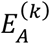 and 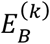 are related by the same isomorphic mappings as 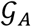 and 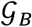, such that:

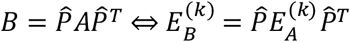

We have 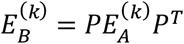 for all 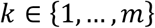. Thus, we have:

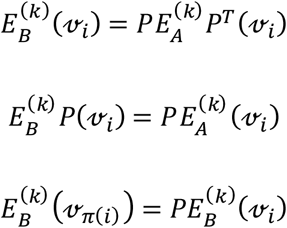

Likewise, we have:

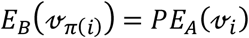

**Lemma 2** demonstrates that the eigenprojection matrices of analogous vertices are related by a row-permutation.

#### Lemma 2.

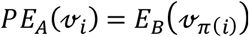.

*Proof*

From Lemma 1 we have:

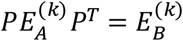

Extracting 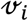 from each eigenspace we have:

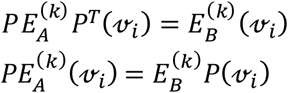

Substituting the vertex permutation, we have:

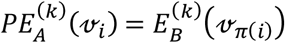

Applying to each eigenspace *k* ∈{1,&, *m*}:

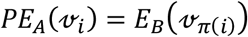

***End of lemma 2***

The linearly permuted relationship between analogous vertices’ eigenprojections allows reframing of the analogy detection task as a linear assignment problem. However, as *P* may be underdefined, it may be not deducible from this equation alone. This issue may be circumvented by applying the row-sorting function *s* to the vertex eigenprojections, neutralising the row-permutation effect of *P*. **Lemma 3** demonstrates that s induces equality between any two row-permutations of a matrix.

#### Lemma 3.

*s*(*X*) = *s*(*PX*) for all 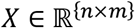 and 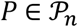.

*Proof*

We have:

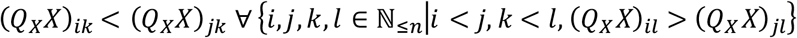

Multiplying internally by *P^T^P*, we have:

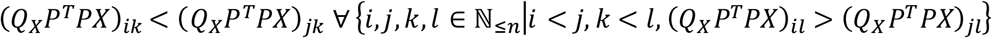

Factorising by (*Q_X_P^T^*), it is evident that *Q_X_P^T^* is a solution to *Q_PX_*. Letting *Q_PX_* = *Q_X_P^T^*, and multiplying from the right by *PX*, we have:

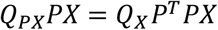

Thus:

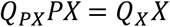

Substituting for the definitions of *s*(*PX*) and *s*(*X*), we have:

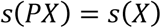

***End of lemma 3***

As analogous vertices eigenprojections are row-permutations of one another, *s* induces equality between them. Applying *s* to a vertex eigenprojection 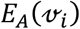, we attain the “ordered vertex eigenprojection”, 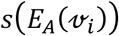. **Lemma 4a** demonstrates that analogous vertices have equal ordered eigenprojections, such that:

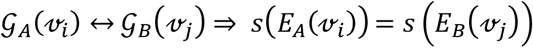

#### Lemma 4a.

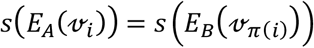.

*Proof*

From Lemma 2 we have:

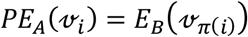

Therefore

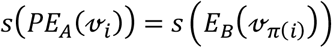

By lemma 3 we may substitute 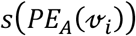 for 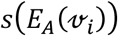.

Thus

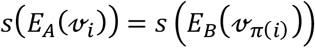

***End of lemma 4a***

**Lemma 4b** demonstrates that if 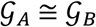, then equality of ordered vertex eigenprojections implies analogy of the corresponding vertices, such that:

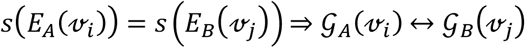

#### Lemma 4b.

If 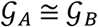 and 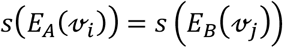 then 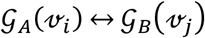.

*Proof*

By lemma 3, there exists some 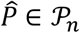 such that:

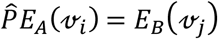

For every *k* ∈ {1,&, *m*} we have:

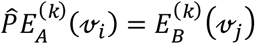

Substituting 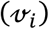 for 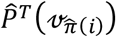, we have:

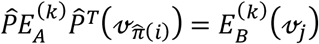

Therefore, for all 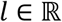 we have

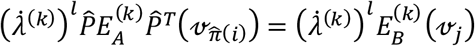

As *A^l^* = *V_A_Λ^l^*(*V_A_*)^*T*^, we have:

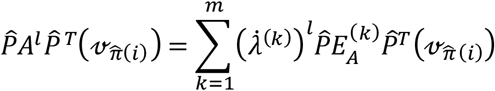

Likewise:

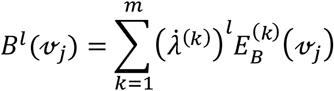

Thus, we have 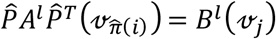 for all 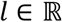. Equal Krylov matrices 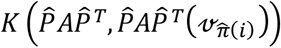 and 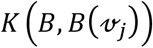 may now be generated such that:

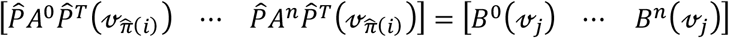

It is noted that these Krylov matrices may be rank deficient if *m* < *n*(Dax, 2017). Deleting the final column, we have:

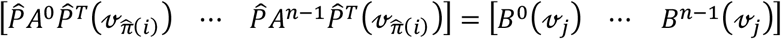

Likewise, deleting the first column we have:

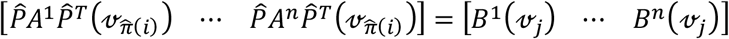

Factorising, we have

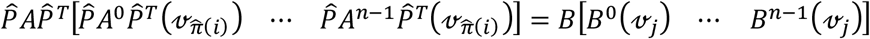

The remaining Krylov submatrices are observed to be equal. Deleting them we observe that 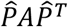 is a solution for *B*. Thus, there exists some 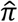 satisfying 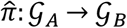 if 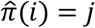, fulfilling the conditions required to demonstrate that 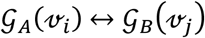. It is noted that the solution to 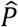 may not be unique, as the Krylov matrices may be rank deficient.

***End of lemma 4b***

Thus, equality of 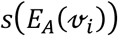 and 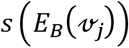 determines analogy of the corresponding vertices, such that:

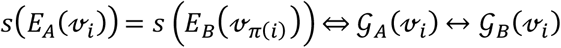

This result implies that the row-ordered vertex eigenprojections may be used to identify analogous vertex pairs, without inferring *P*. Thus, the vertex analogy testing problem is linearised exactly, providing a polynomial time route to extract an isomorphic mapping 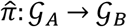.

At this point, a critical distinction from Klus’ method must be noted (Klus and Sahai, 2018). Klus sorted the columns of 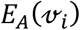 separately, such that the ordered vertex eigenprojection was given by:

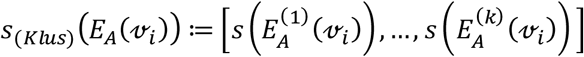

Consequently, different permutations may be applied to each column, so that the vertex eigenprojection and its ordered image may not be related by a linear permutation. Clearly, Klus’ ordering transformation maps analogous vertices to equal ordered vertex eigenprojections, such that:

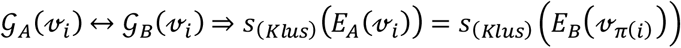

However, the converse does not hold, as Klus’ method may also map some non-analogous vertices to the same ordered vertex eigenprojection if the columns of their eigenprojections can be sorted equally. Therefore:

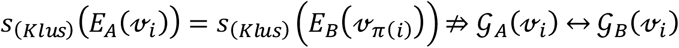

We now return to the description of our method. As 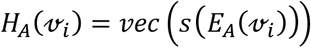, equality of 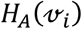 and 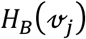 may also be employed to test analogy of 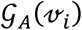 and 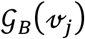. By applying a second row-ordering transformation to *H_A_* and *H_B_* it is observed that they are brought into a matrix form, *s*(*H_A_*), which is invariant to graph permutation. **Lemma 5** demonstrates that *s*(*H_A_*) is a permutation invariant representation of 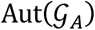.

#### Lemma 5.

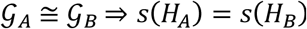.

From lemmas 4a and 4b we have:

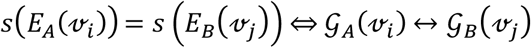

As 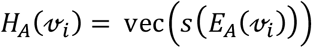 and 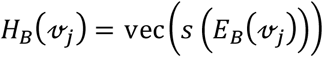, we have:

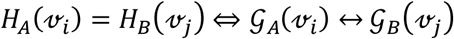

Thus

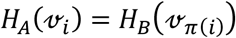

Therefore *PH_A_* = *H_B_* if 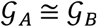. From lemma 3 we have *s*(*PH_A_*) = *s*(*H_A_*). Therefore

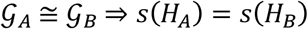

Therefore, application of the row-ordering function *s* to *H_A_* returns a such that *s*(*H_A_*) = *s*(*H_B_*) if 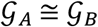.

***End of lemma 5***

### Constructing the isomorphic mapping

Now that individual vertex analogies may be identified, the problem of constructing the isomorphic mapping from individual analogies is addressed. A precursor to the permutation matrix, 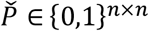, may be generated by testing each vertex pair for analogy, such that:

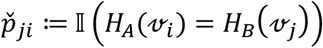

Equivalently,

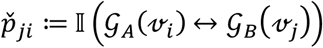

If the solution to *π* is unique, each vertex has only one analogy and 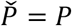. However, if multiple isomorphisms exist, some vertices will have multiple analogies and 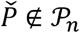. To resolve this issue, analogy-preserving perturbations must be applied to *A* and *B* to induce uniqueness to the isomorphic mapping. Let 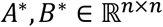 denote the perturbed matrices. Let 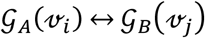 *A* and *B* may be perturbed at *a_ii_* and *b_jj_* to generate isomorphic matrices *A*\* and *B*\* with a unique isomorphism 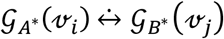.

Let 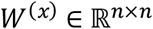 such that:

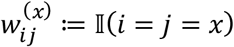

The perturbed matrices *A*\* and *B*\* are declared such that:

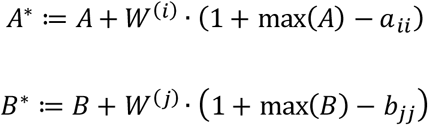

Thus:

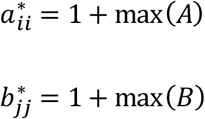

Thus, characteristically weighted self-loops are added to analogous vertices, thereby inducing a unique analogy between them. The perturbation scheme is depicted graphically in **Figure 1**.

**Figure 1.**
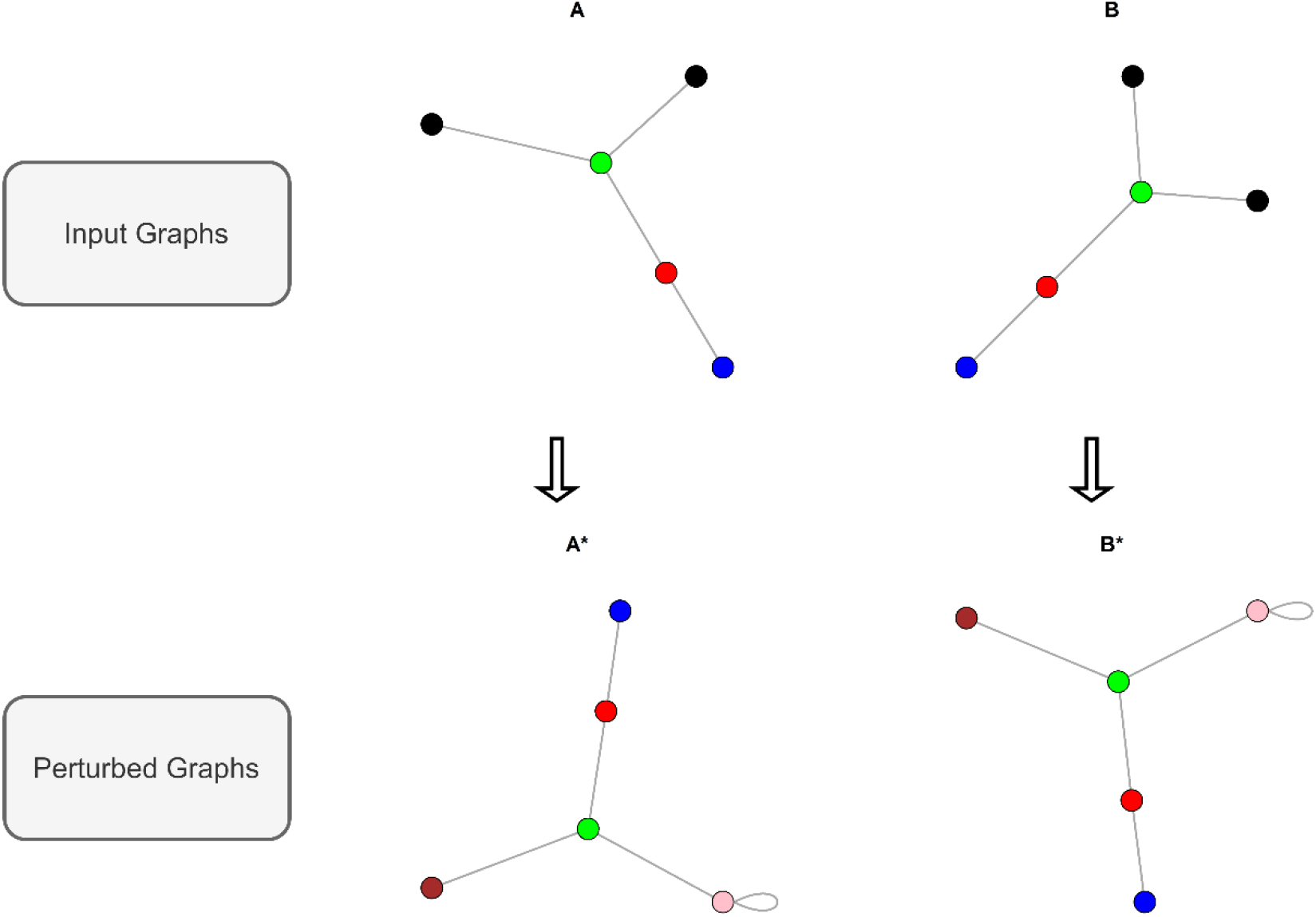
Graphical illustration of analogy testing and perturbation in an isomorphic tuple of degenerate graphs. A and B represent input graphs, and vertex colours indicate their analogy to one another. As multiple analogies exist between the black vertices, perturbation is required. A* and B* demonstrate the perturbed graphs in which self-loops have been added to one of the black vertice in each graph, removing any instances of multiple analogy.

**Lemma 6** demonstrates that perturbation of analogous vertices 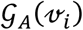 and 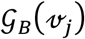 induces a unique analogy 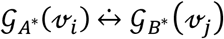. Thus, if 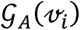 and 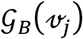 had other analogies, these are broken in the perturbed graphs. Thus, perturbation may be targeted to break graph symmetries and remove degeneracy.

#### Lemma 6.

Let 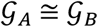. Let *A*\* and *B*\* denote perturbations of *A* and *B* such that:

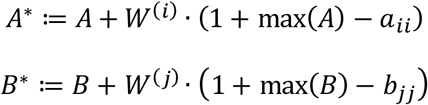

If 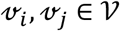 such that 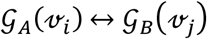 then 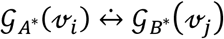.

*Proof*

We have max(*A*) = max(*B*). Hence:

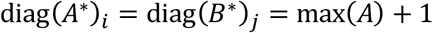

Hence, for all 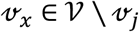 we have:

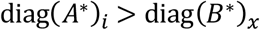

Therefore 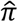 cannot satisfy 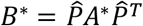 unless 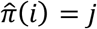. Therefore 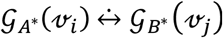.

***End of lemma 6***

**Lemma 7** demonstrates that if 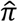 satisfies the perturbed equality 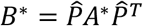 then it satisfies the original equality 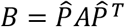. Thus, if 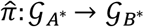 then 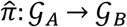. Consequently, the search for 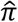 may continue in the perturbed graphs.

#### Lemma 7.

If 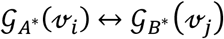 and 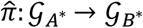 then 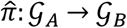.

We have:

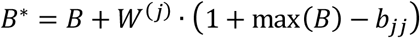

Factorising the left side of the equality we have:

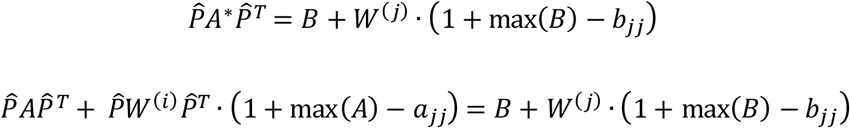

As 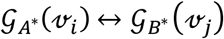 we can substitute 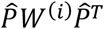 for *W*^(*j*)^, thus:

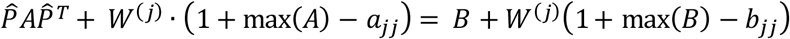

As max(*A*) = max(*B*) and *a_ii_* = *b_jj_*, we have:

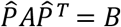

Therefore 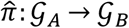.

***End of lemma 7***

Here, a subtle distinction from Klus’ perturbation strategy is noted. Klus also perturbed graphs through the addition of self-loops, which were weighted 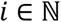 at the *i*th iteration. This strategy does not guarantee induction of a unique analogy as identical self-loops may already exist on other vertices. By weighting the new self-loop as (1 + max(*A*)) uniqueness is guaranteed.

If a non-uniquely analogous vertex pair is perturbed in this way, a unique analogy is induced and the number of isomorphisms is thereby reduced. Vertex analogies may now be checked in the perturbed matrices *A*\* and *B*\*, such that:

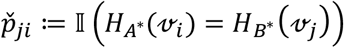

If 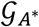 and 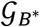 remain degenerate, other non-uniquely analogous vertex pairs may be sought in the perturbed graphs. Thus, perturbations may be applied iteratively, such that:

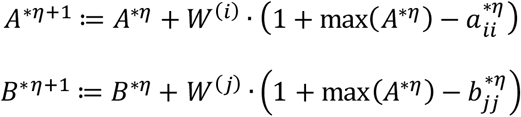

where *A*^\**η*^ and *B*^\**η*^ denote the graphs after 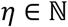 perturbation iterations and 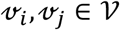 such that 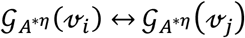. Naively, analogies could be sought, and perturbations applied for each vertex in turn. After *n* – 1 iterative perturbations, each entry on the perturbed matrix diagonals would be distinct, implying that any solution to 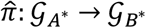 would be unique. However, if some vertices already have unique analogies, these need not be perturbed. Thus, fewer than *n* iterations are required to induce a unique isomorphism between 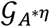 and 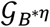. This procedure constitutes **Algorithm 1**, a naïve approach which is included here to aid understanding.

Now the task of graph canonical labelling is addressed. From lemma 1 it is known that *E_A_* is invariant to *A*. Likewise, for all 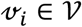 the ordered vertex eigenprojections *H_A_*(*v_i_*) remain invariant across the permutation group of 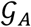. Thus, vertices 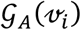 and 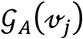 may be ordered canonically by 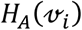 and 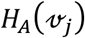 if 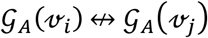. However, analogous vertices cannot be ordered canonically in this manner, as 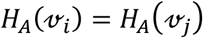 iff 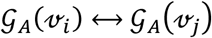. Therefore, using the ordered eigenprojections, vertices may only be ordered up to analogy. This result is consistent with our definition of analogy, which implies equivalence of analogous vertices under permutation. Hence, uniqueness of the canonical ordering requires that no analogous vertices exist within the graph. Thus, a perturbation strategy must be implemented to order vertices canonically. This is achieved by partially ordering the vertices according to the row-ordering of *H_A_*, then applying a perturbation to the first vertex in the ordering with an analogy. It is observed that, as analogous vertices are identified by identical rows in *H_A_*, that any non-random row-ordering operation will assign adjacent order positions to analogous vertices. Thus, it is guaranteed that no analogy will be missed if only rows with adjacent order positions are tested. This procedure constitutes **Algorithm 2**, which dominates Algorithm 1 in terms of computational efficiency, as fewer analogy tests are required.

### Algorithms

#### Algorithm 1

In algorithm 1, the ordered vertex eigenprojection matrices are extracted for both graphs and every vertex pair is compared. If vertices have multiple analogies, one of the non-uniquely analogous pairs is selected and a characteristic perturbation applied to both vertices. This concludes the first iteration. Ordered vertex eigenprojection matrices may now be extracted from the perturbed graphs and further perturbation are applied in this manner until no vertices remain with multiple analogies.

##### Algorithm 1 pseudocode

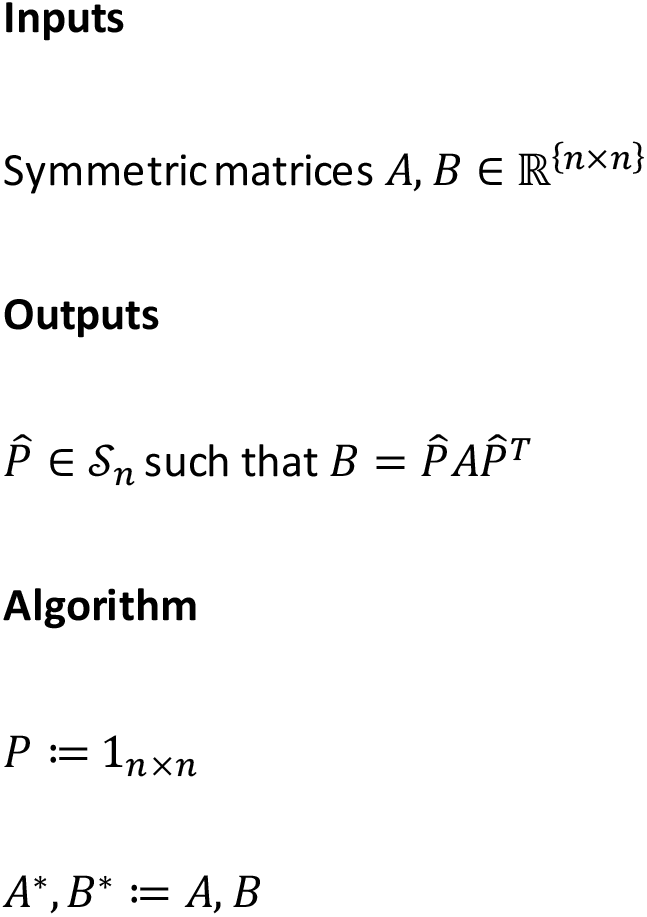
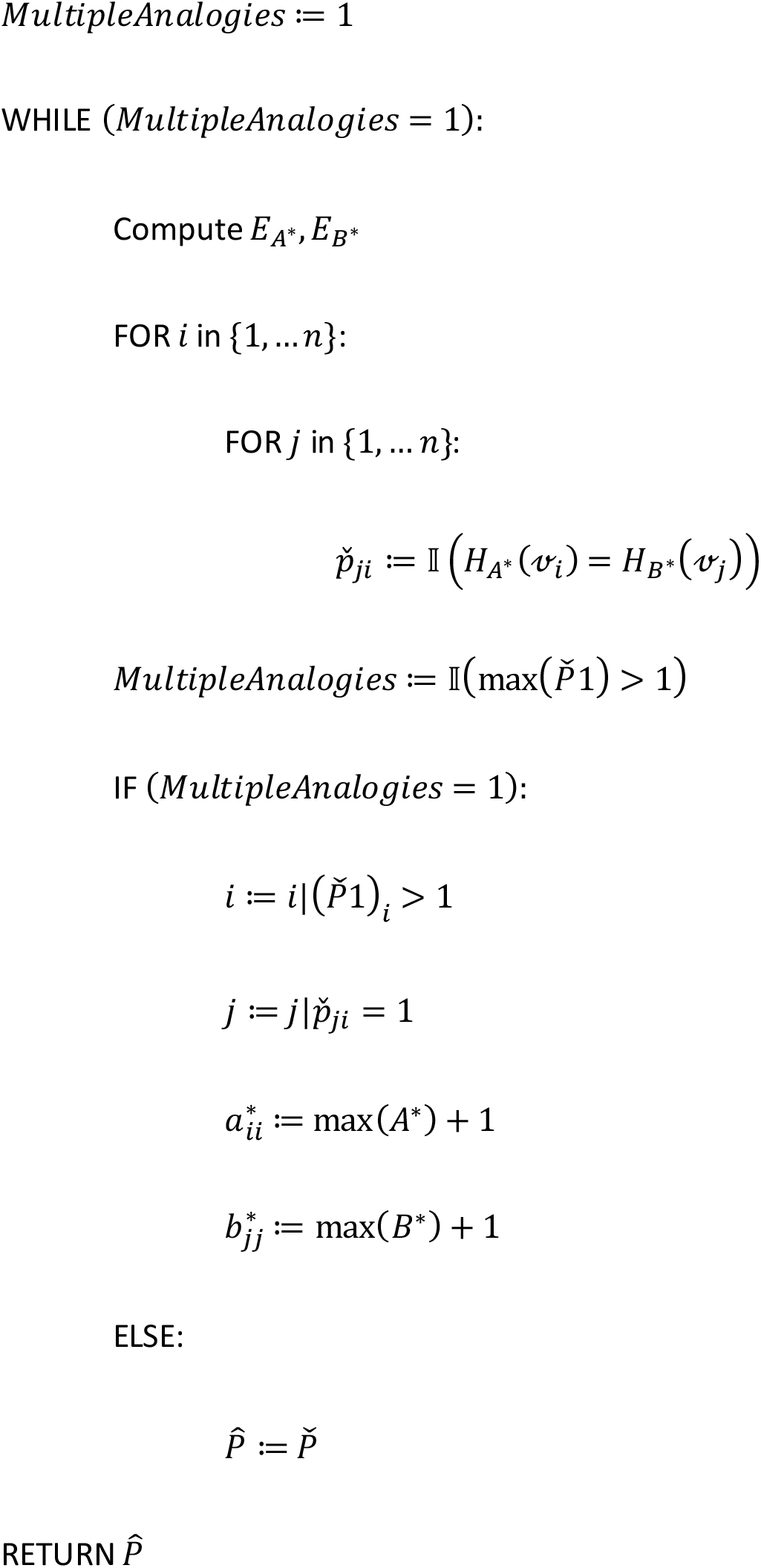

#### Algorithm 2

In algorithm 2, canonical labellings are inferred for each graph separately. For each graph, the ordered vertex eigenprojections are extracted. Then, vertices themselves are ordered according to their ordered vertex eigenprojections. This results in a partial ordering, such that non-analogous pairs are ordered whilst analogous pairs remain unordered. Following the ordering, the first *n* – 1 vertices are tested for analogy with the subsequent vertex. The first vertex identified to have an analogy is perturbed by adding a characteristically weighted self-loop. This concludes the first iteration. Ordered vertex eigenprojections are extracted now from the perturbed graph and further perturbations applied until no vertex analogies remain.

##### Algorithm 2 pseudocode

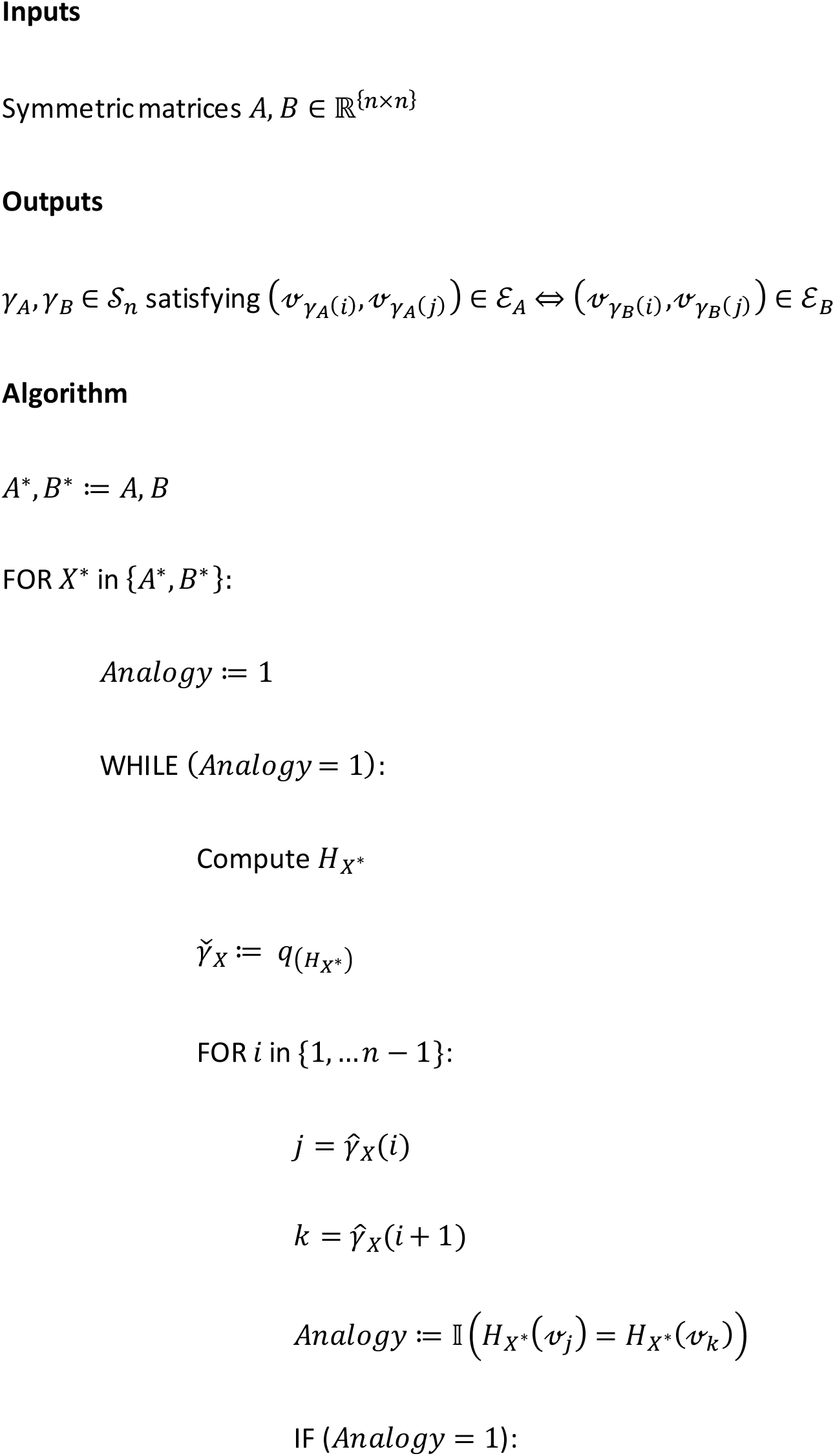
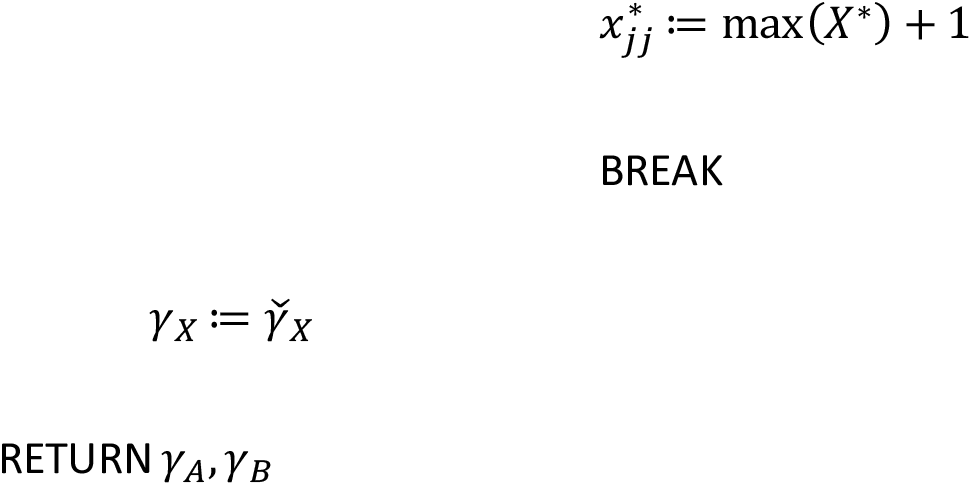

### Implementation

The performance of algorithm 2 was tested on real-word and synthetic graphs. Real-world graphs were extracted from a public repository of brain networks (Amunts *et al*., 2013; Rossi and Ahmed, 2015). As regularly structured synthetic graphs present some of the most challenging problems for traditional algorithms (Babai *et al*., 1982), a 100-vertex ring graph and Johnson graphs (Holton and Sheehan, 1993) with parameters (n=9, k=4) and (n=10, k=3) were also included. In each experiment, a permutation was applied to one of the graphs to generating a tuple of isomorphic graphs. The algorithm was subsequently applied to infer an isomorphic mapping. Computation was performed with R and RStudio using iGraph and abind libraries (R Core Team, 2021; RStudio Team, 2021; Csardi and Nepusz, 2006; Plate and Heiburger, 2016). Algorithm 2 was employed to infer isomorphic mappings. To minimise discrepancies between matrix eigenvalues due to numerical fuzz, the presumption of isomorphism was invoked, and eigenvalues of both matrices were averaged. This procedure is permissible if isomorphism is formally tested later through validation of the isomorphic mapping. A numerical tolerance of 10^-10^ was applied when testing equality of eigenvalues and eigenvectors, respectively. All code required to reproduce the findings of this study is provided at github.com/robertoshea/graph_isomorphism. The BLISS canonical labelling algorithm (Junttila and Kaski, 2007) was included for comparison, via iGraph. Experiments were conducted on a standard laptop with 16Gb RAM and an Intel Core i7 CPU.

## Results

Algorithm 2 identified a correct isomorphic mapping in each experiment (**Table 1**). No experiment required more than *n* perturbations. The longest running experiment “bn_mouse_visual_cortex_2”, with 193 vertices and 23 distinct eigenvalues, required 159 iterations over 245 seconds to infer an isomorphic mapping. The eigenprojection ordering algorithm inferred correct isomorphic mappings in each experiment. BLISS failed to infer correct mappings in any of the experiments involving degenerate graphs.

**Table 1.** Experiment characteristics and results.

| Graph | Vertices | Distinct eigenvalues | Edges | Algorithm | Correct solution | Iterations | Time (secs) |
| --- | --- | --- | --- | --- | --- | --- | --- |
| cat mixed species brain 1 | 65 | 65 | 1139 | Eigenprojection ordering | TRUE | 1 | 0.25 |
| cat mixed species brain 1 | 65 | 65 | 1139 | BLISS | TRUE | NA | 0 |
| macaque rhesus brain 1 | 242 | 234 | 4090 | Eigenprojection ordering | TRUE | 9 | 47.62 |
| macaque rhesus brain 1 | 242 | 234 | 4090 | BLISS | FALSE | NA | 0.01 |
| macaque rhesus brain 2 | 91 | 23 | 628 | Eigenprojection ordering | TRUE | 33 | 5.8 |
| macaque<br>rhesus<br>brain 2 | 91 | 23 | 628 | BLISS | FALSE | NA | 0 |
| macaque<br>rhesus<br>cerebral<br>cortex 1 | 91 | 59 | 1615 | Eigenprojection<br>ordering | TRUE | 1 | 0.3 |
| macaque<br>rhesus<br>cerebral<br>cortex 1 | 91 | 59 | 1615 | BLISS | TRUE | NA | 0 |
| macaque<br>rhesus<br>interarea<br>I cortical<br>network<br>2 | 93 | 6 | 2667 | Eigenprojection<br>ordering | TRUE | 91 | 18.55 |
| macaque<br>rhesus<br>interarea<br>I cortical<br>network<br>2 | 93 | 6 | 2667 | BLISS | FALSE | NA | 0.03 |
| mouse<br>brain 1 | 213 | 213 | 21807 | Eigenprojection<br>ordering | TRUE | 1 | 4.86 |
| mouse<br>brain 1 | 213 | 213 | 21807 | BLISS | TRUE | NA | 0.02 |
| mouse<br>visual<br>cortex 1 | 29 | 23 | 44 | Eigenprojection<br>ordering | TRUE | 5 | 0.15 |
| mouse<br>visual<br>cortex 1 | 29 | 23 | 44 | BLISS | FALSE | NA | 0 |
| mouse<br>visual<br>cortex 2 | 193 | 23 | 214 | Eigenprojection<br>ordering | TRUE | 159 | 267.29 |
| mouse<br>visual<br>cortex 2 | 193 | 23 | 214 | BLISS | FALSE | NA | 0.01 |
| ring 100 | 100 | 51 | 100 | Eigenprojection<br>ordering | TRUE | 3 | 1.22 |
| ring 100 | 100 | 51 | 100 | BLISS | FALSE | NA | 0 |
| johnson<br>9 4 | 126 | 5 | 1260 | Eigenprojection<br>ordering | TRUE | 7 | 0.95 |
| johnson<br>9 4 | 126 | 5 | 1260 | BLISS | FALSE | NA | 0.01 |
| johnson<br>10 3 | 120 | 4 | 1260 | Eigenprojection<br>ordering | TRUE | 7 | 0.78 |
| johnson<br>10 3 | 120 | 4 | 1260 | BLISS | FALSE | NA | 0 |

Table 1. Experiment characteristics and results. In each experiment a random permutation was applied to the input graph and each algorithm was applied to infer an isomorphic mappings. Algorithm 2 was employed for Eigenprojection Ordering. BLISS iterations were not provided by the implementation used.

## Discussion

This analysis unifies the problems of graph canonisation and isomorphism testing via the tensor of eigenprojections. It is demonstrated that the ordered vertex eigenprojections provide a signature by which vertex analogy may be tested. By reshaping the ordered vertex eigenprojections into a matrix structure and applying a second ordering, a permutation invariant graph representation *s*(*H_A_*) is achieved. This result is consistent with the quadratic nature of the problem – that two ordering operations should be required to recover the equivalence broken by two permutation operations. It is conjectured that this representation is unique to 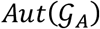, and is therefore a canonical form. This hypothesis is supported by the observation that equality of *s*(*H_A_*) and *s*(*H_B_*) implies that every vertex in 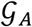 has an analogy in 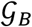 and vice versa. Correctness of this hypothesis would imply that *s*(*H_A_*) is invariant over 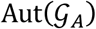 and unique to 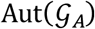, fulfilling the requirements of a canonical representation. Therefore, if this conjecture was proven correct, isomorphism testing could proceed in 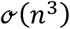 time through equality testing of *s*(*H_A_*) and *s*(*H_B_*). Analysis of this conjecture is left for further research.

This study demonstrates that a graph with analogous vertices may be labelled canonically through repeated iterations of partial ordering and perturbation. As *n* such iterations may be required, this algorithm guarantees canonical vertex labelling in 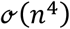 time. Algorithm 2 dominates algorithm 1, by ordering vertices such that analogous vertices are assigned adjacent positions in the ordering. Consequently, analogy tests are only required between vertices with adjacent positions in the ordering. Thus, *n* – 1 analogy tests are required to demonstrate uniqueness of each vertex in algorithm 2, compared with *n*^2^ – *n* in algorithm 1.

The algorithms described in this paper assume that the graph is undirected. If the graph is directed, the adjacency matrix may be asymmetric and uniqueness of the tensor of eigenprojections is not guaranteed. The extension of these algorithms to directed graphs is left for further research. The algorithms do not require any spectral conditions, such as positive definiteness or bounded eigenvalue multiplicity. Where multiple isomorphic mappings exist, a single solution to 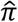 is returned, however the full set of solutions may also be computed by changing the order of perturbations.

As the algorithm tests equality of eigenvalues and elements of eigenprojections, numerical stability issues are expected (Klus and Sahai, 2018). This is a critical limitation of the above algorithms to practical graph isomorphism problems. In some cases, numerical stability may be achieved through appropriate rounding or numerical tolerance. However, errors are likely in large graphs with similar eigenvalues. To assess the suitability of the algorithm for a particular use case it is recommended to perform simulations by duplicating one of the graphs, permuting it and attempting to recover an isomorphic mapping. This procedure may also be used to optimise numerical tolerance settings.

The algorithm presented here demonstrates that any problem which may be reduced to a graph isomorphism problem may be solved in polynomial time. Further research will investigate developments to the algorithm to solve the subgraph isomorphism problem and the general case-quadratic assignment problem in polynomial time.

## Acknowledgements

The author would like to acknowledge the advice of Reimer Kuhn and Stefan Izaak at the Department of Theoretical Physics, King’s College London. The author would like to acknowledge the support of Vicky Goh and Gary Cook at the Department of Cancer Imaging, King’s College London; and Sophia Tsoka at the Department of Informatics, King’s College London.

## Conflicts of Interest

The author has no conflicts of interest to declare.

## Availability of Code and Materials

All code required to reproduce the results of this analysis is implemented in the R language and provided at github.com/robertoshea/graph_isomorphism.

## Funding

Authors acknowledge funding support from the UK Research& Innovation London Medical Imaging and Artificial Intelligence Centre; Wellcome/Engineering and Physical Sciences Research Council Centre for Medical Engineering at King’s College London (WT 203148/Z/16/Z); National Institute for Health Research Biomedical Research Centre at Guy’s& St Thomas’ Hospitals and King’s College London; Cancer Research UK National Cancer Imaging Translational Accelerator (A27066).

## Notes

https://github.com/robertoshea/graph_isomorphism

### Competing Interest Statement

The authors have declared no competing interest.

### Summary of Updates

Lemma 4b has been revised to correct an error, and prove the proposition via Krylov matrices.

